# Transcriptome based changes in expression of detoxification genes of *Bemisia tabaci* under destruxin A treatment

**DOI:** 10.1101/2023.06.28.545844

**Authors:** Can Zhang, Jianling Guo, Shaukat Ali, Bao-Li Qiu

## Abstract

**Background:** *Bemisia tabaci* is an important agricultural pest that has been causing significant economic losses to crops across the globe. Destruxins are secondary metabolites of entomopathogenic fungi which can be used as a potential biopesticide against *B. tabaci*. However, little is known about the molecular mechanism regulating the defense response of *B. tabaci* post destruxin application.

**Results:** In this study, we explored the molecular responses of *B. tabaci* exposed to destruxin A (DA) using RNA-Seq and differentially expressed gene (DEG) analysis. A total of 1702, 616, and 555 DEGs were identified in *B. tabaci* after 4, 8, and 12 h of destruxin A treatment. In addition, 40 putative detoxification-related DEGs, including 29 cytochrome P450s (P450s), 5 glutathione S-transferases (GSTs), and 5 carboxylesterases (CarEs) were also identified. Quantitative real-time PCR analysis indicated that the expression profiles of 19 random DEGs were consistent with the RNA-Seq results.

**Conclusion:** These findings serve as valuable information for a better understanding of the interaction and molecular mechanisms involved in the defense response of *B. tabaci* against DA.

## Background

The whitefly, *Bemisia tabaci* (Gennadius) (Hemiptera: Aleyrodidae), is a serious pest of crops in different regions of the world (Barbosa *et al*., 2014; De Barro *et al*., 2011). Direct damage of *B. tabaci* occurs as a result of feeding plant sap from the phloem, excreting honeydew which results in ‘sooty molds’ on leaves, causing adverse effects on crop productivity (Oliveira *et al*., 2001; Perring *et al*., 2018). Moreover, *B. tabaci* can transmit more than 350 plant viruses to commercial crops (Polston *et al*., 2014; Götz *et al*., 2016; Lu *et al*., 2019). Being the most notorious invasive species of this complex, *B. tabaci* Middle East-Asia Minor 1 (MEAM1) has a wide range of host plants (Perring *et al*., 2018; Shadmany *et al*., 2019). The control of MEAM1 *B. tabaci* is often based on conventional insecticides including carbamates, organophosphates, and pyrethroids, which has resulted in the development of insecticide resistance (Horowitz *et al*., 2011; Shah *et al*., 2020; Abubakar *et al*., 2022). Also, it causes pollution to the environment and affects human health.

Destruxins, secondary metabolites produced by entomopathogenic fungi like *Metarhizium anisopliae*, can potentially be used for various purposes like V-ATPase inhibitors and biocontrol agents (Liu and Tzeng, 2012). About 39 derivatives of destruxins have been identified to date (Liu and Tzeng, 2012). Among them, destruxin A (DA) has received much attention because of its high insecticidal activity (Pedras *et al*., 2002). The application of DA to control *B. tabaci* can slow down its pest resistance and have lower negative impacts on its natural enemy (Hu *et al*., 2009).

However, studies on the insecticidal mechanism of DA have been performed against insects including *Drosophila melanogaster, Spodoptera litura, Plutella xylostella*, and *Bombyx mori* (Pal *et al*., 2007; Han *et al*., 2013; Meng *et al*., 2013; Wang *et al*., 2019; Wang *et al*., 2020; Yin *et al*., 2022). A few reports are available on the insecticidal mechanism DA against whitefly (Hu *et al*., 2009; Zhang *et al*., 2017). As a xenobiotic, how DA overcomes the detoxification and metabolism responses of *B. tabaci* needs further research.

This study reports the results from the sequencing and assembly of the transcriptomes of two groups (with and without DA treatment) of *B. tabaci*. The readings were first assembled through de novo genome assembly. DEGs were detected and functionally annotated, and the gene expression patterns for the selected DEGs were validated by qRT-PCR. Our data will provide insights into the molecular mechanisms of the detoxification response of *B. tabaci* under DA treatment.

## Materials and Methods

### Insect rearing

MEAM1 whiteflies were collected from cotton plants and were reared at South China Agricultural University (SCAU), in greenhouse facilities on *Gossypium hirsutum* (Malvaceae), Lu-Mian 32 at 26±1 °C, 70±10% R.H., and 14:10 h light/dark photoperiod.

### Destruxin A treatments

Destruxin A was extracted and purified from strain MZ16 of *Metarhizium anisopliae* isolated from MeiZhou, Guangdong. DA was diluted to 1 mg/ml with DEPC water and was stored at −20 °C untill further use.

The median lethal concentration (LC_50_) of DA was determined in a pilot experiment, in which *B. tabaci* adults were fed on six increasing doses of DA solution with 15% sucrose and 5% yeast powder. The dose that came closest to killing 50% of the adults (17.80 μg/mL) within 24 h was then selected for experiments.

After filtering and sterilizing with a 0.22 μm filter membrane, the solution was transferred to an artificial feeding device (transparent double-way tube) and covered with parafilm. 100 pairs of 3-day-old whitefly adults were transferred into the device from the other side and sealed with pierced parafilm. The feeding devices were wrapped in the lower half with black cloth and placed in a light incubator (26 ± 1°C, relative humidity 55 ± 10%, photoperiod L: D = 14 h: l0 h). After feeding for 4 h, 8 h, and 12 h, alive adults from each treatment were collected and stored at -80 °C. The control was fed by the feeding solution without DA, and each treatment was repeated 3 times.

### RNA extraction and transcriptome sequencing

The alive *B. tabaci* adults were used for RNA extraction. Total RNA was extracted using Trizol reagent (Magen Biotechnology Co., Ltd, China), and the integrity, purity, and concentration of the RNA were verified by Agilent 2100 Bioanalyzer (Agilent, United States).

After the RNA had been completely extracted, mRNA was purified from the total RNA using poly-T oligo-attached magnetic beads. Then, the enriched mRNA was fragmented into short fragments using a fragmentation buffer. To synthesize first- and second-strand cDNAs, random hexamers, dNTPs, and DNA polymerase I were used. The double-stranded cDNA was purified with magnetic beads following the ligation of fragments with sequencing adaptors enriched by PCR amplification. To qualify and quantify the sample libraries, Agilent 2100 Bioanalyzer and ABI Step One Plus Real-Time PCR System were used following the sequencing on the Illumina HiSeqTM4000 system (Illumina, United States). Illumina sequencing was performed at the Beijing Genomics Institute (BGI-Shenzhen, China).

### Genome mapping and analysis of differentially expressed genes

Clean reads were obtained after removing raw reads with adapters and unknown bases by SOAPnuke software. After filtration by Bowtie 2 (Langmead *et al*., 2012), clean reads were mapped with the reference gene and reference genome by HISAT (Kim *et al*., 2015). Finally, all data were normalized as fragments per kilobase of transcript per million fragments mapped (FPKM). Differential expression analysis was carried out by a strict approach, and the threshold p-value was determined by using the false discovery rate (FDR) methodology for analyzing multiple tests (Kim and van de Wiel, 2008). A standard threshold (FDR ≤ 0.001 and log2 ratio≧1) was set to identify significantly differentially expressed genes (DEGs) in the libraries.

### Functional analysis of differentially expressed genes

The differential genes were functionally mapped to the Gene Ontology (GO) database and Kyoto Encyclopedia of Genes and Genomes (KEGG) pathways. The *Bemisia tabaci* genome (Chen *et al*., 2016) was set as the background. The corrected p-value < 0.05 and functions with FDR ≤ 0.01 was set as a threshold for the enrichment of significantly different functions.

### Validation of DEG libraries by quantitative real-time PCR (qRT-PCR)

Eighteen different genes were randomly selected from the detection results and verified by qRT-PCR of MEAM1 *B. tabaci*. The primers were shown in Table 1 and EF1α was selected as the internal reference gene. The cDNA (template) was obtained by using PrimeScript™ RT Reagent Kit (TaKaRa, Japan). The qRT-PCR reaction includes 10 μL SYBR qPCRMix (TOYOBO, Japan), 0.3 μM each primer, 7.5 μL sterilized water, and 2μL cDNA. The program was as follows: 95°C for 3 min, 40 cycles of 95 °C for 15 s, 60 °C for 30s, and 72 °C for 45s. The 2^-ΔΔCt^ technique was used to calculate relative expression levels using β-actin as an internal control (Wang *et al*., 2009). Three biological replicates were performed for each treatment, and three technical replicates were performed for each sample. All the experiments were performed with the Bio-Rad CFX-96 system.

**Table 1.**
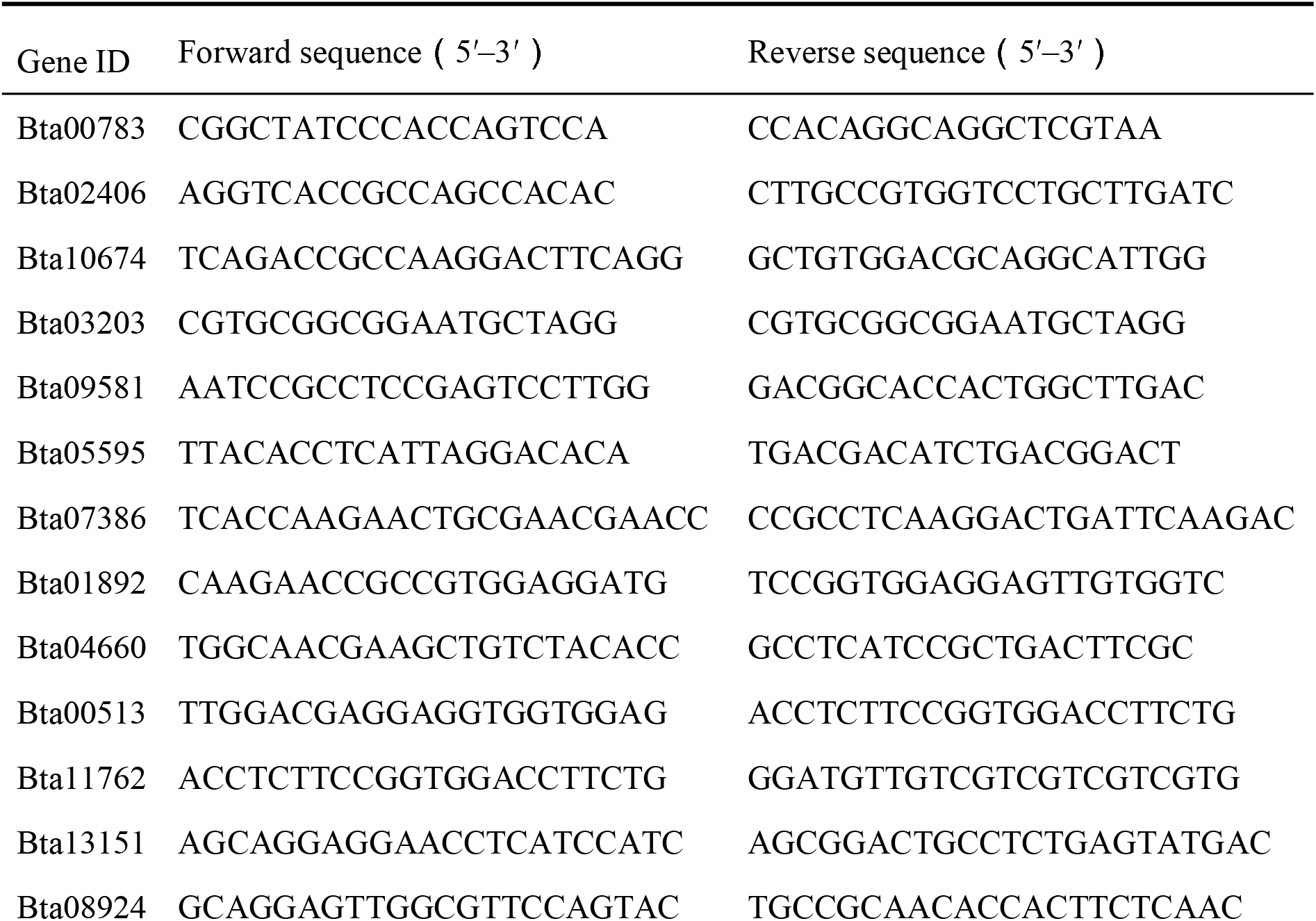
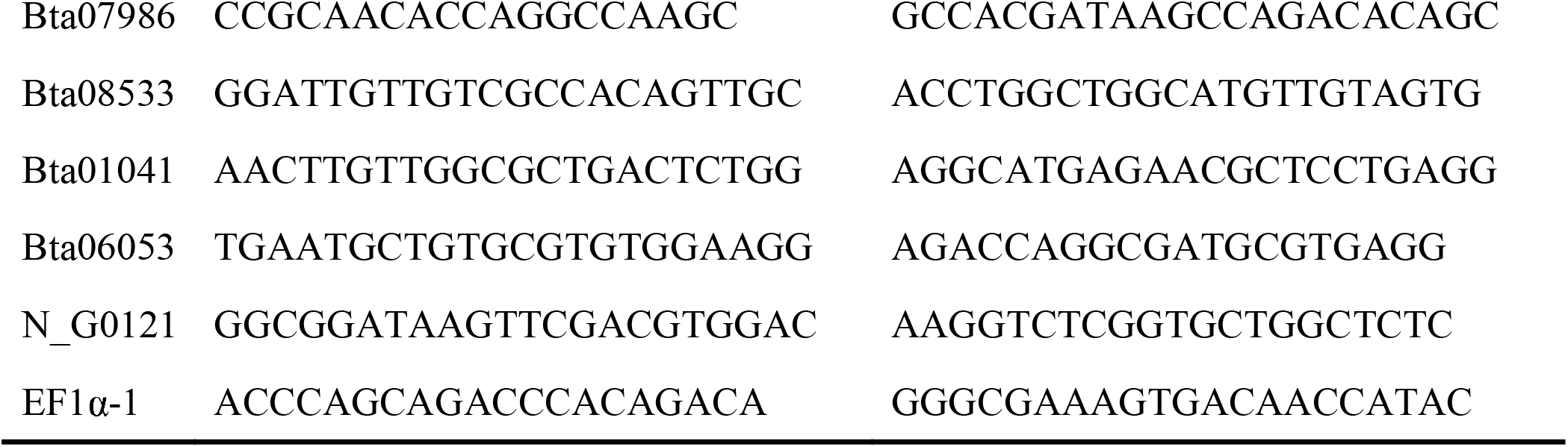
Primers used in qRT-PCR.

## Results

### Summary of sequencing and mapping to reference genome

The results of the sequencing analysis of experimental insects under different temperature treatments are shown in Supplementary table-1. The Q20 values were all over 99%; the Q30 value was in the range of 96.78∼97.78%. This indicates that the construction quality of the sequencing library is good, and the data obtained by sequencing are accurate and reliable. Among the data of the clean read libraries, 76.01% to 82.02% of the clean reads were successfully mapped to the reference genome.

### Dynamics of DA Responsive DEGs

The transcripts of the whiteflies were treated in two groups: not treated with destruxin A as a whole (CK), and treated with destruxin A as a whole (DA). The Venn diagram showed that there are 814 genes expressed specifically in the control group, 684 genes expressed specifically in the DA group, and a total of 17,688 genes expressed in both the control and treated groups (Fig. 1A).

**Fig. 1.**
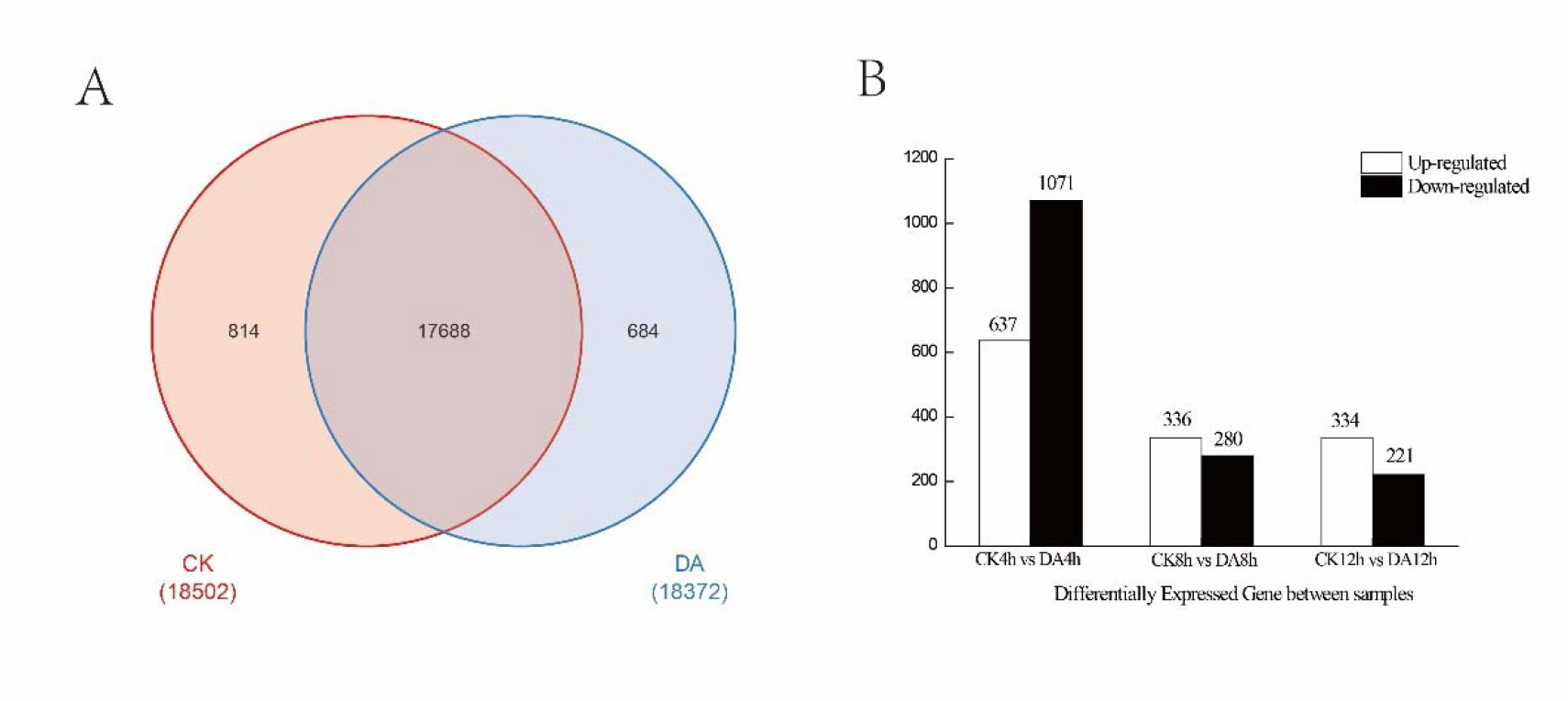
Dynamics of DA responsive DEGs. A. Venn diagram analysis; B. DEGs after treated by destruxin A. FDR ≤ 0.001, log_2_ (fold change) ≥ |1.0|. Note: CK (not treated by destruxin A), DA (treated by destruxin A)

The PossionDis method was used for differential gene expression analysis and the differentially expressed genes between the treatment and control groups of whiteflies treated with DA for 4, 8, and 12 hours were identified through statistical analysis. The transcriptional expression pattern of relevant genes was determined in adult whiteflies before and after treatment with DA. The results showed that the transcriptional regulation of a large number of genes was rapidly activated in the transcriptome of whiteflies after treatment with DA. Compared to the control group, genes with expression differences of more than two-fold were considered as differentially expressed genes. A total of 637 genes with up-regulated expression and 1071 genes with down-regulated expression were screened at 4 hours, 336 genes with up-regulated expression and 280 genes with down-regulated expression were screened at 8 hours, and 334 genes with up-regulated expression and 221 genes with down-regulated expression were screened at 12 hours (Fig. 1B). Differentially expressed genes between CK and DA are shown in Table S2.

#### 3.3 Functional Annotation of DGEs

Based on the differential gene detection results before and after treatment with DA, a comprehensive analysis was carried out to identify DA-responsive genes in whiteflies by combining GO and KEGG annotation results. As shown in Fig. 2, a total of 1092 differential genes were annotated in the GO enrichment, which accounted for 37.96% of all differential genes.

**Fig. 2.**
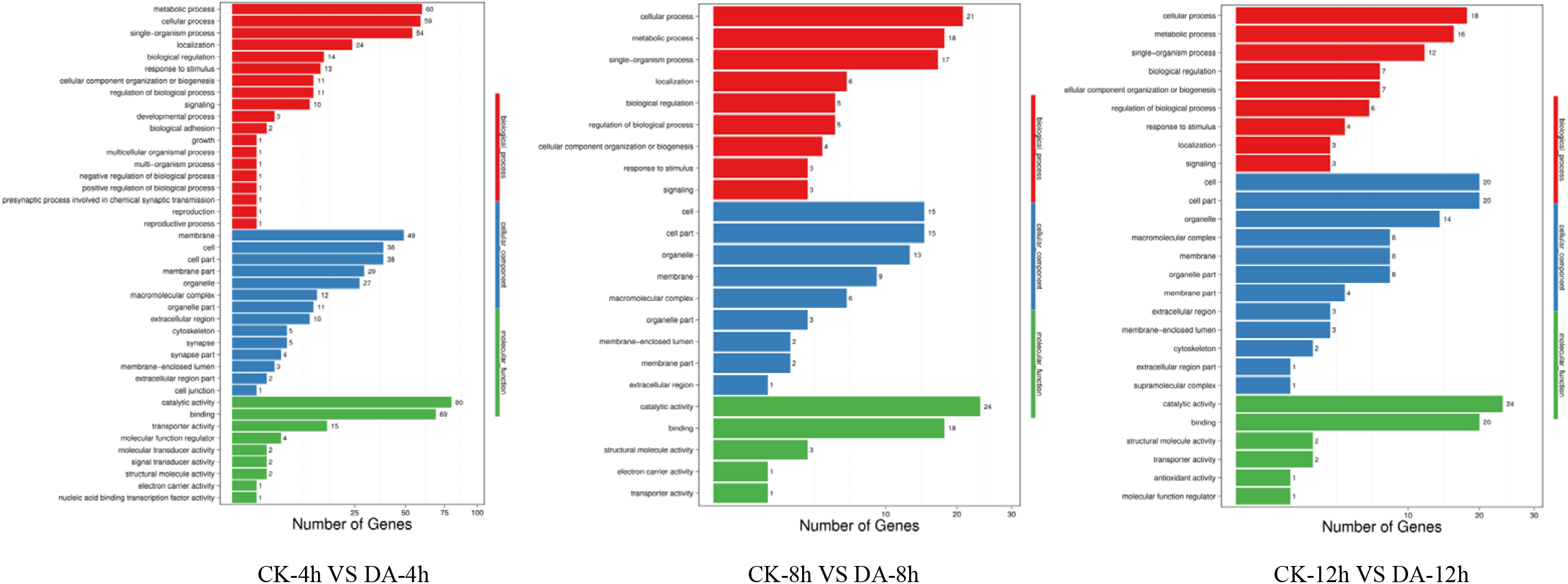
GO classification of DEGs in *Bemisia tabaci* Note: The X axis represents the number of DEG. The Y axis represents the GO term.

To further analyze the functions of all significantly differentially expressed genes in their corresponding pathways, GO enrichment and KEGG pathway analyses were performed. A total of 1708 deferentially expressed genes were identified after 4 hours of treatment, of which 679 genes were annotated with GO functions. Catalytic activity was the most enriched term (11.78%) following binding (10.16%), metabolic processes, and cellular processes. After 8 and 12 hours of treatment, the most enriched terms were also involved in catalytic activity, binding, cellular processes, and metabolic processes (Fig. 3). The KEGG enrichment pathways of highly expressed genes in *B. tabaci* treated by destruxin A were shown in Table 2.

**Table 2.** The main KEGG enrichment pathways of highly expressed genes in *Bemisia tabaci* treated with destruxin A at different time intervals

| KEGG categories | KEGG pathways | Gene numbers |  |  |
| --- | --- | --- | --- | --- |
|  |  | 4 h | 8 h | 12 h |
| Organismal Systems: Immune system | Chemokine signaling pathway | 30 | 44 | 19 |
| Human Diseases: Infectious diseases: Parasitic | Amoebiasis | 39 | 7 | 8 |
| Metabolism: global and overview maps | Carbon metabolism | 37 | 7 | 5 |
| Human Diseases: Infectious diseases: Bacterial | Vibrio cholerae infection | 36 | 3 | 7 |
| Cellular Processes: Cell growth and death | Apoptosis | 33 | 12 | 7 |
| Cellular Processes: Cellular community - eukaryotes | Tight junction | 32 | 8 | 9 |
| Organismal Systems: Circulatory system | Adrenergic signaling in cardiomyocytes | 28 | 6 | 3 |
| Human Diseases: Cancers: Overview | Transcriptional misregulation in cancer | 26 | 10 | 12 |
| Organismal Systems: Immune system | Fc gamma R-mediated phagocytosis | 25 | 38 | 15 |
| Cellular Processes: Cell growth and death | Cell cycle | 26 | 5 | 9 |
| Environmental Information Processing: Signal transduction | Hippo signaling pathway - fly | 26 | 7 | 7 |
| Metabolism: Carbohydrate metabolism | Amino sugar and nucleotide sugar metabolism | 23 | 7 | 1 |
| Human Diseases: Cardiovascular diseases | Dilated cardiomyopathy | 23 | 8 | 3 |
| Cellular Processes: Cell growth and death | Oocyte meiosis | 23 | 4 | 7 |
| Organismal Systems: Endocrine system | Oxytocin signaling pathway | 20 | 5 | 3 |
| Metabolism: Carbohydrate metabolism | Pyruvate metabolism | 19 | 2 | 3 |
| Organismal Systems: Endocrine system | Estrogen signaling pathway | 16 | 4 | 3 |
| Organismal Systems: Circulatory system | Cardiac muscle contraction | 15 | 4 | 1 |
| Organismal Systems: Immune system | Toll and Imd signaling pathway | 13 | 6 | 11 |
| Environmental Information Processing: Signal transduction | TGF-beta signaling pathway | 7 | 2 | 7 |

**Fig. 3.**
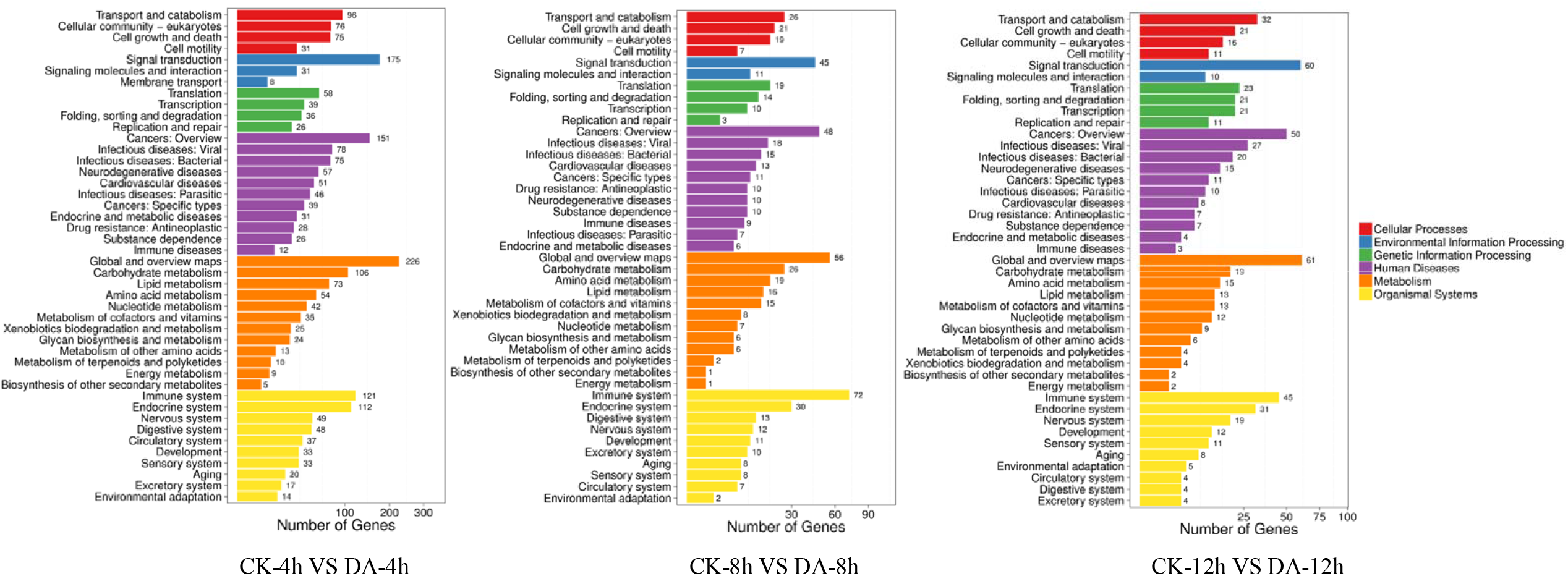
Pathway classification of DEGs in *Bemisia tabaci*. Note: The X axis represents the number of DEG. The Y axis represents the functional classification of KEGG

### DEGs involved in detoxification

A total of 39 detoxification genes showed significant differences in expression in response to DA, including 29 cytochrome P450 (CYP450) genes, 5 Glutathione S-transferase (GST) genes, and 5 Carboxylesterase (CarE) genes, as shown in Table 3. Most of the CYP450 genes were down-regulated in response to DA after being treated with DA for 4 hours; for example, *CYP6DW4, CYP4CT1*, and *CYP402C11* showed a downregulated expression of 8.37-fold, 7.36-fold, and 7.34-fold, respectively. Among the GST genes, one gene was up-regulated by 7.43-fold, and 3 genes were down-regulated by 1.43, 4.13, and 9.93-fold, respectively. Two Carboxylesterase genes, *Carboxylesterase* and *Esterase 6* exhibited upregulated expression at 4 h after treatment, with 7.35 and 7.17 fold.

**Table 3.**
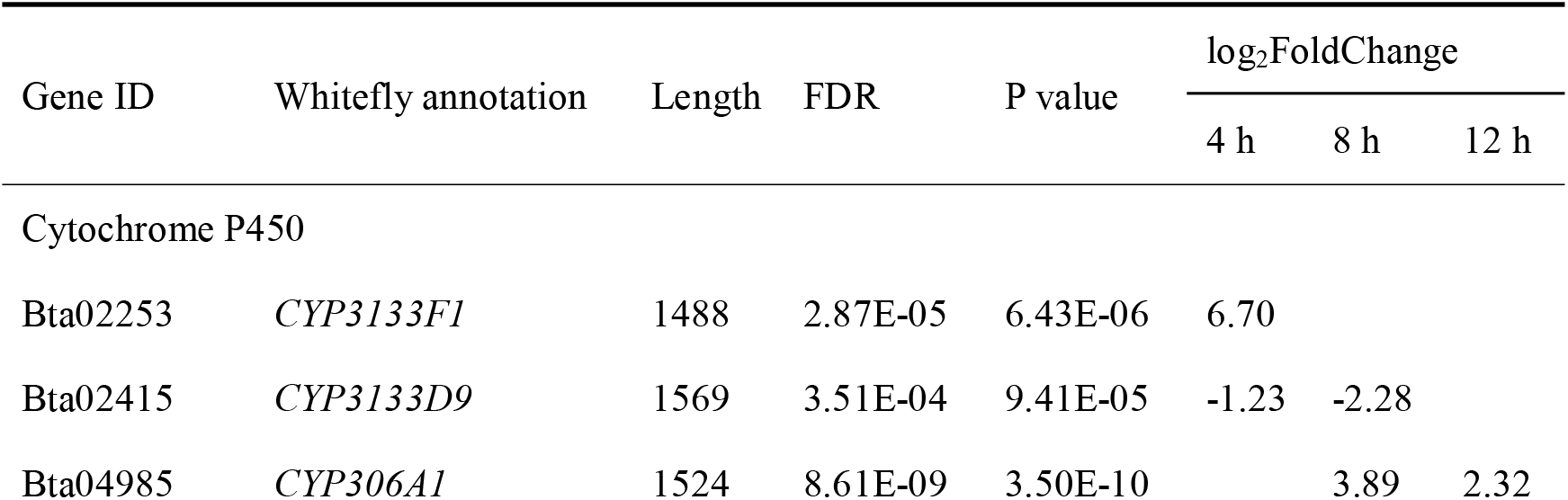
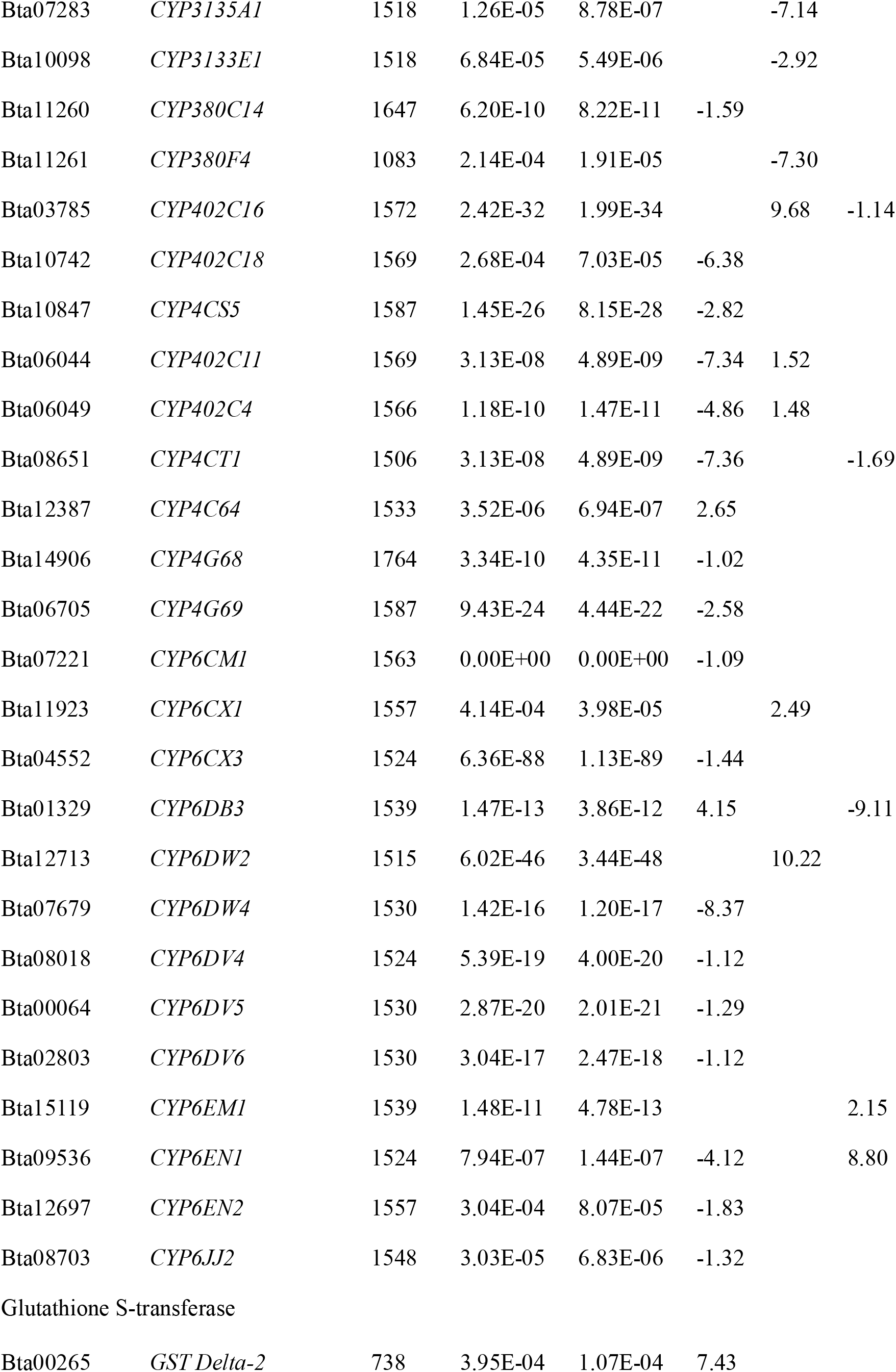
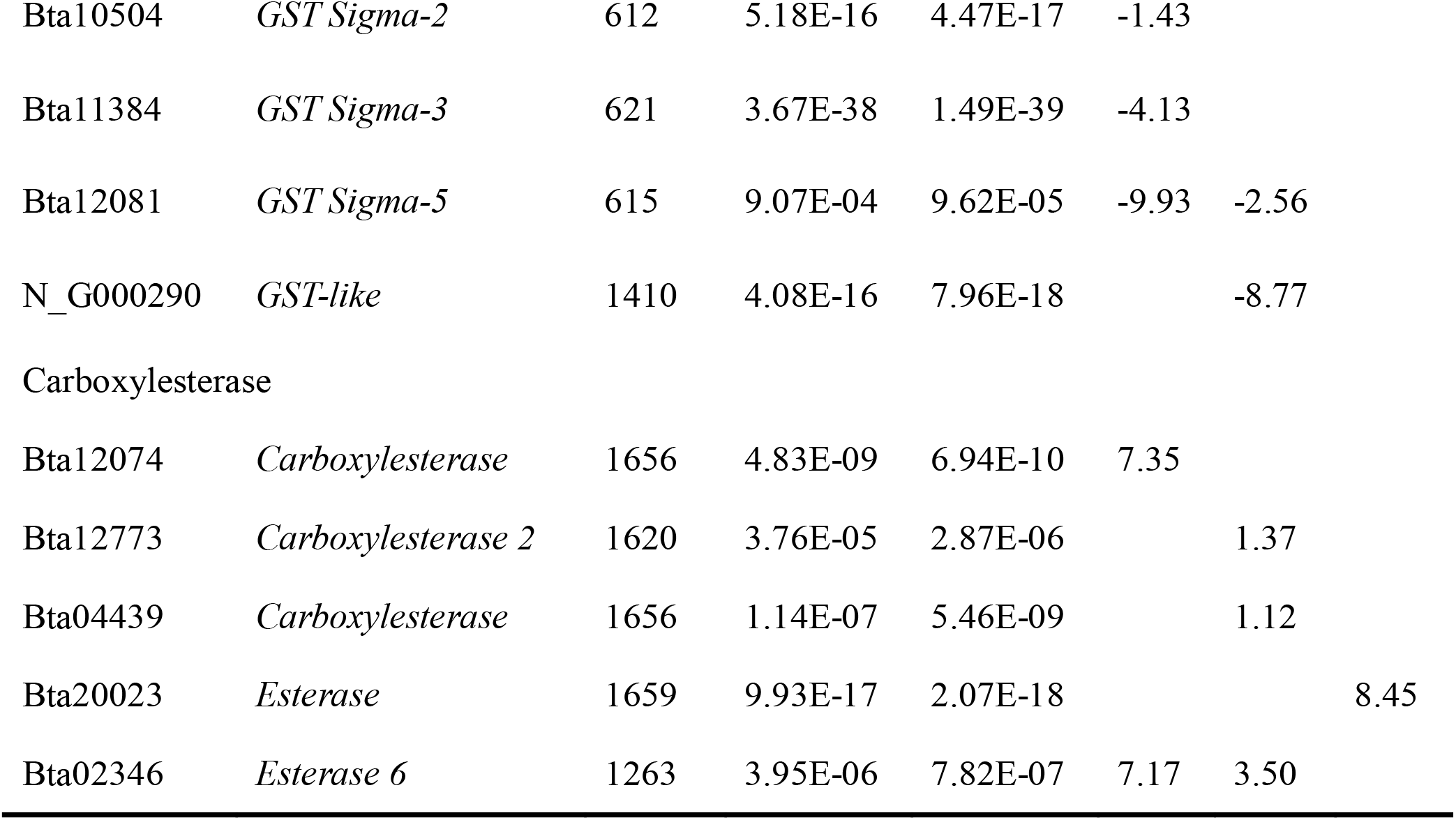
Differential expressed genes of detoxification in *Bemisia tabaci* after treated with destruxin A

After 8 hours of DA treatment, 7 CYP450 genes were up-regulated, with the highest up-regulation being 10.22-fold (*CYP6DW2*), and 4 CYP450 genes were down-regulated, with the lowest down-regulation being 7.30-fold (*CYP380F4*). Two GST genes, *GST-like*, and *GST Sigma-5* showed a down-regulated expression of 8.77 and 2.56-fold. After 8 h, *Carboxylesterase 2, Carboxylesterase, and Esterase 6* genes were up-regulated by 1.37-, 1.12- and 3.5-fold, respectively.

After 12 hours of DA treatment, *CYP6EN1, CYP306A1*,and *CYP6EM1* genes were up-regulated by 8.80-2.32-, 2.15-fold; and 3 CYP450 genes were down-regulated, with the lowest down-regulation being 9.11-fold (*CYP6DB3*). The *esterase* gene was up-regulated by 8.45-fold after 12 h.

### qRT-PCR validation of differentially expressed genes

To verify the accuracy of the differential gene expression analysis data from transcriptome, we randomly selected 18 differentially expressed genes and validated their expression in the treatment and control groups using qRT-PCR. The cDNA from the treatment and control groups of the whiteflies were used as templates, and the *EF1*α gene was used as an internal reference. The results showed that the qRT-PCR results of these genes were consistent with the transcriptome analysis (Fig. 4), indicating the stability and reliability of the sequencing data.

**Fig. 4.**
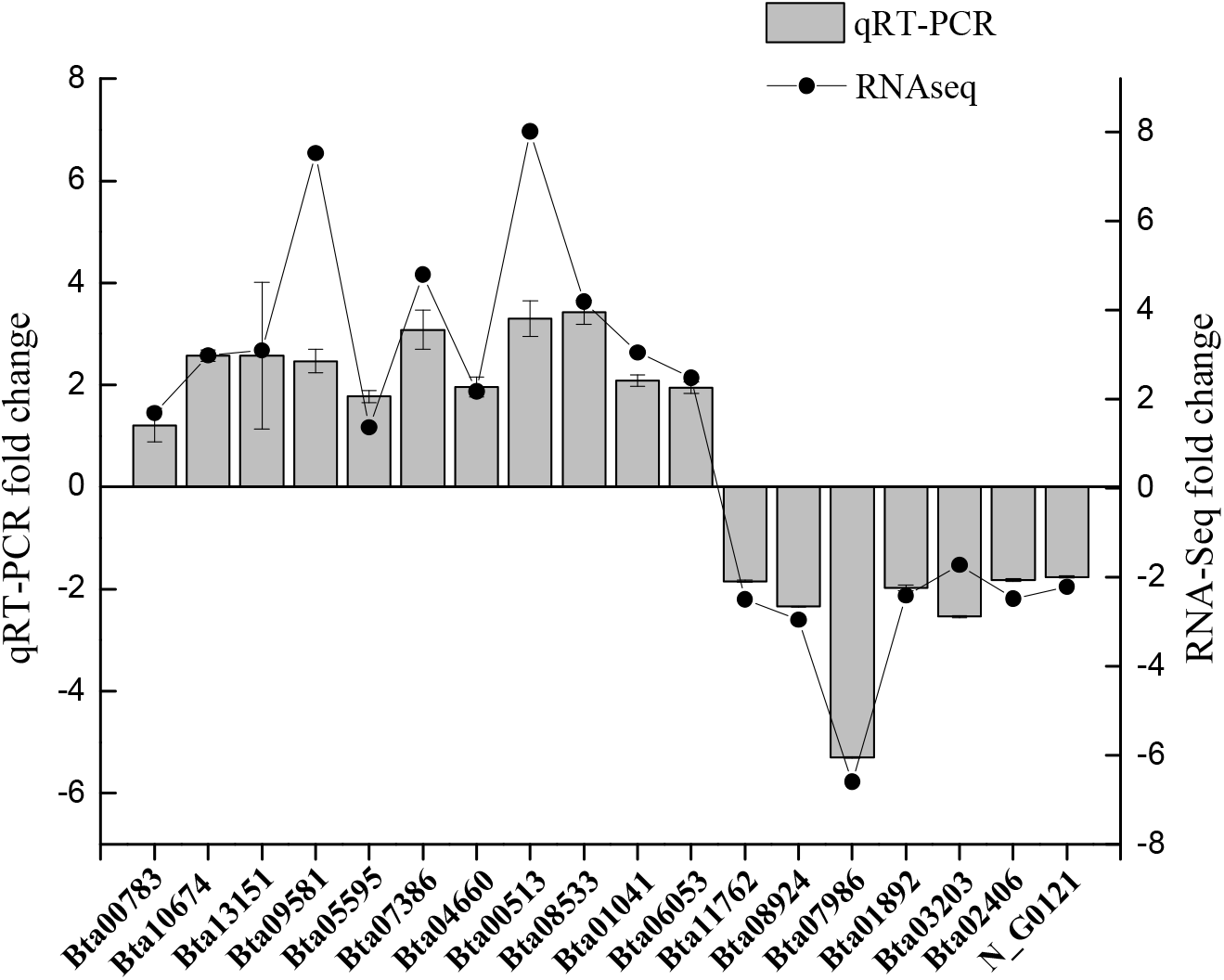
qRT-PCR validation of differently expressed genes. Bars represent standrad error of means based on three replicates.

## Discussion

As a secondary metabolite of entomopathogenic fungi, destruxins have low toxicity to non-target animals (Strasser *et al*., 2000), transient storage in plants, and a rapid degradation process (Rios-Moreno *et al*., 2016). The use of destruxin A as bio insecticide is considered a promising biological alternative to control MEAM1 *B. tabaci*. In this study, we sequenced and analyzed the transcriptome of the whiteflies treated with the DA at LC_50_ concentration using the Illumina sequencing platform, obtaining transcriptomes of treated samples at 4 h, 8 h, and 12 h and control samples. After removing adapter sequences and low-quality reads, we obtained an average data volume of 5.65 Gb per sample, indicating successful library construction and high-quality sequencing.

Functional annotation of DEGs revealed their involvement in transportation and metabolism, signal transduction, cancer, detoxification of exogenous substances, and the immune system, indicating that DA can affect multiple pathways in MEAM1 *B. tabaci*. Further analysis showed that these DEGs were mainly enriched in pathways related to chemokine signaling, apoptosis, and carbon metabolism, as well as pathogen infection and the immune system, cellular processes, and metabolism, indicating that insecticide A has a significant impact on the immune system and metabolism of MEAM1 *B. tabaci*. These results are consistent with previous studies on the effects of DA in *S. litura* and *B. mori* (Han *et al*., 2013; Gong *et al*., 2014; Wang *et al*., 2023a). DA seems to have multi-targets and affect multiple life processes of insects.

Dysfunction of detoxification and metabolism was an important factor that lead to insecticides poisoning. As a mycotoxin secreted by entomopathogenic fungi, DA needs to overcome the insect defense system to reach the target organ. Insects’ detoxifying enzyme system was affected by DA based on digital gene expression profiles (Han *et al*., 2013; Gong *et al*., 2014). Cytochrome P450, a polygenic enzyme superfamily, plays important roles in growth, development, nutrition, and detoxification of xenobiotics in insects (Feyereisen, 2011; Lu *et al*., 2021). Cytochrome P450 is the main detoxifying enzyme system responsible for detoxification, which is critical for the metabolism of diverse exogenous and endogenous compounds, including pesticides in insects (Iga and Kataoka, 2012; Liu *et al*., 2015). In our study, 29 CYP450 genes of MEAM1 *B. tabaci* showed up or down regulation, especially during the early stage of treatment (4h), which indicated roles of cytochrome P450s resisting to DA and meantime the insect could resist DA by CYP450s. In the supergene family of cytochrome P450, genes of CYP4 and CYP6 family are considered to be more involved in insecticide resistance than other cytochrome P450 families (Scott, 2008; Zhu *et al*., 2016). Studies have shown that *CYP6CM1, CYP6CX1*, and CYP4C64 genes are closely related to imidacloprid and thiamethoxam resistance in the whitefly species complex (Karunker *et al*., 2008; Zhuang *et al*., 2011; Barman *et al*., 2022; Guo *et al*., 2023). Of the 21 differentially regulated CYP450 genes after 4 h treatment, 18 genes were downregulated, indicating that DA may inhibit detoxification by suppressing cytochrome P450 genes so as to increase the attack on *B. tabaci*.

GSTs belong to a multigene family of dimeric multifunctional proteins that play important roles in the detoxification of heterogeneous compounds including drugs, herbicides, and insecticides (Hayes *et al*., 2005). Insects obtained resistance through GSTs by metabolizing or sequestering pesticides or by preventing pesticide-induced oxidative stress (Pavlidi *et al*., 2018). Enhanced expression of GSTs has been shown to be a mechanism of resistance to DDT and organophosphates, and is also associated with resistance to pyrethroids in some insects (Huang *et al*., 1998; Lumjuan *et al*., 2005; Ranson *et al*., 2001). In our results, only one GST gene was up-regulated, and all other genes were down-regulated, which indicated the inhibitory effect of DA on GST in *B. tabaci*. Carboxylesterase plays an important role in the metabolic resistance in many insects, especially in the detoxification of organophosphorus, carbamates, and pyrethroids (Yan *et al*., 2009). In organophosphate-resistant MEAM1 *B. tabaci*, CarE activity was significantly increased compared with sensitive strains, and CarE1 and CarE showed higher expression levels (Rauch e*t al*., 2003; Alon *et al*., 2008). In this study, unlike CYP450 and GST, CarE showed an activation after DA treatment, indicating that CarE could still perform detoxification function after DA treatment. This may be due to the mutual coordination of the detoxification function in *B. tabaci* after cytochrome P450 and GST were inhibited. As a global pest, MEAM1 *B. tabaci* has shown increased insecticide resistance, reflecting its strong detoxification ability. By analyzing gene expression profiles from transcriptomes, we gain a better understanding of underlying molecular mechanisms involved in the toxic response to DA in MEAM1 *B. tabaci*. This knowledge will provide foundation in developing new applications for entomopathogenic fungi toxins and in advancing research on new insecticides for pest control.

## Conclusions

In conclusion, this study has generated comprehensive transcriptomes of *B. tabaci* to destruxin A at different time points, and qRT-PCR experiments showed that the results of transcriptome analysis are reliable. The genes differentially expressed in the controls and DA-treated were largely identified and functionally annotated. DA treatment led to a rapid response of the detoxification of *B. tabaci*. Our results provide a rich data resource for an in-depth study of the potential targets of DA and elucidation of the mechanism of action of DA as a biopesticide.

## Abbreviations

DA: destruxin A
*B. tabaci*: *Bemisia tabaci*
*S. Litura*: *Spodoptera litura*
*B. Mori*: *Bombyx mori*
LC_50_: the median lethal concentration
%: Percent
CYP450: cytochrome P450
GST: Glutathione S-transferase
CarE: Carboxylesterase

## Author contributions

CZ and JG conducted the experiment. CZ wrote the manuscript. SA and BQ reviewed this article. All authors read and approved the final manuscript.

## Ethics declarations

### Ethics approval and consent to participate

Not applicable.

### Consent for publication

Informed consent was obtained from all individual participants included in the study.

