## Supplementary Table 1 for "Transcriptome based changes in expression of detoxification genes of *Bemisia tabaci* under destruxin A treatment"

Supplementary table -1: Transcriptome sequencing statistics

| Sample | Total Raw Reads(Mb) | Total Clean Reads(Mb) | Total Clean Bases(Gb) | Clean Reads Q20(%) | Clean Reads Q30(%) | Clean Reads Ratio(%) | Total Mapping Ratio |
| --- | --- | --- | --- | --- | --- | --- | --- |
| CK_4 h | 59.04 | 56.09 | 5.61 | 99.11 | 96.78 | 95.00 | 82.02% |
| DA_4 h | 59.04 | 56.51 | 5.65 | 99.33 | 97.72 | 95.71 | 80.25% |
| CK_8 h | 59.04 | 56.47 | 5.65 | 99.31 | 97.72 | 95.65 | 76.01% |
| DA_8 h | 59.04 | 56.60 | 5.66 | 99.31 | 97.78 | 95.86 | 77.34% |
| CK_12 h | 59.04 | 56.55 | 5.66 | 99.33 | 97.77 | 95.79 | 78.42% |
| DA_12 h | 59.04 | 56.54 | 5.65 | 99.31 | 97.73 | 95.77 | 79.72% |
